# OxyR regulates the oxidative stress response in *Zymomonas mobilis* during oxic growth and anoxic biofuel fermentation

**DOI:** 10.64898/2026.05.23.727408

**Authors:** Emma C. Boismer, Magdalena M. Felczak, Kevin S. Myers, Michaela A. TerAvest

**Affiliations:** Department of Biochemistry and Molecular Biology, Michigan State University, East Lansing, MI, USA, 48824; Great Lakes Bioenergy Research Center; Wisconsin Energy Institute, University of Wisconsin-Madison, Madison, Wisconsin, USA, 53726

## Abstract

The bacterium *Zymomonas mobilis* is widely studied for its potential as an industrial biofuel producer. Anoxic fermentation by *Z. mobilis* in lignocellulosic hydrolysate can generate bioethanol from renewable plant biomass. In this study, we deleted a gene from the *Z. mobilis* genome encoding a homolog of OxyR, a transcription factor that activates an oxidative stress response in bacteria to reduce reactive oxygen species (ROS). Deletion of this transcription factor inhibited growth of *Z. mobilis* in oxic, but not anoxic, conditions in laboratory media. A ROS probe revealed that the *oxyR* response is required to reduce intracellular ROS during oxic growth. Importantly for biofuel production, the absence of *oxyR* inhibited growth and delayed ethanol production during anoxic hydrolysate fermentation. To determine the source of oxidative stress in hydrolysate, we grew *ΔoxyR* in a synthetic hydrolysate containing known inhibitors found in hydrolysate. There was no growth defect in *ΔoxyR* in the synthetic hydrolysate, indicating that known inhibitory compounds are not the source of anoxic oxidative stress. We determined that Ammonia-Fiber Expansion (AFEX) switchgrass hydrolysate contains significant peroxide concentrations. Addition of catalase to hydrolysate improves growth of both *ΔoxyR* and wild-type *Z. mobilis* in hydrolysate. This study uncovers an important source of stress to *Z. mobilis* during biofuel fermentation.

**Importance:** Fermentation of non-food biomass is a promising avenue for sustainable production of fuels and chemicals, but several challenges currently limit applicability of this technology. One major hurdle is that when biomass is deconstructed into a fermentable form, many byproducts are generated that inhibit microbial fermentation. Here, we investigated how a fermentative bacterium, *Zymomonas mobilis*, experiences oxidative stress during anoxic biomass fermentation, and identified genes important in this response. These findings provide a better understanding of the stresses faced by *Z. mobilis* during biofuel production. Fully understanding the effects of hydrolysate on biofuel-producing microbes is crucial for optimizing production and making carbon-neutral fuel a reality.

## Introduction

Lignocellulosic biomass is one of the most abundant, renewable, and inexpensive resources on Earth and may be an important feedstock for production of fuels and chemicals (1). Lignocellulosic biomass processing typically involves harvest, pretreatment with solvents or ammonia to break down lignocellulosic polymers, and enzymatic hydrolysis to produce a liquid called lignocellulosic hydrolysate (2). This hydrolysate can be fermented using bacteria or yeast to convert sugars into fuels or other products. *Zymomonas mobilis* is a top microbial candidate for bioethanol production from lignocellulosic biomass, due to its capacity for converting glucose to ethanol with up to 97% efficiency under anoxic conditions (3). However, the ethanol yield is lower when *Z. mobilis* is exposed to oxygen and side products are generated, including acetaldehyde, acetate, and acetoin (4–6).

Poor oxic growth in *Z. mobilis* is partially attributed to acetaldehyde accumulation, which occurs because electrons from NAD(P)H are transferred to the respiratory electron transport chain (ETC) instead of reducing acetaldehyde to ethanol (5). To mitigate acetaldehyde build-up, *Z. mobilis* oxidizes it to acetate, using acetaldehyde dehydrogenase (5). Much work has been done to characterize the *Z. mobilis* ETC (7–13) and the *Z. mobilis* oxygen response has been studied through transcriptomics, metabolomics, proteomics, and CRISPR interference (CRISPRi) (6, 14, 15). *Z. mobilis* upregulates ETC components in response to oxygen, likely to increase aerobic respiration for oxygen removal (14). Iron-sulfur (FeS) cluster maintenance and biogenesis proteins are also upregulated aerobically, which may play a role in mitigating oxidative damage to FeS cluster containing enzymes (14). It has recently been discovered that respiration is essential for *Z. mobilis* oxic growth, for the purpose of eliminating oxygen rather than energy generation via oxidative phosphorylation (16). Despite recent advancements in understanding the *Z. mobilis* response to oxygen, the transcriptional regulation of this response remains unknown.

One transcription factor that may contribute to the oxygen response is OxyR, a conserved bacterial transcription factor that acts as a biosensor of reactive oxygen species (ROS) through oxidation of its critical cysteine residues. OxyR is a LysR-type regulator, containing a DNA-binding domain and a ligand-binding domain (17). Homologs of OxyR exist across the bacterial phyla of *Proteobacteria*, *Bacteroidetes*, and *Actinobacteria* (18). The mechanism by which OxyR senses oxidative stress to regulate gene expression has been well-studied in *E. coli*. Hydrogen peroxide (H_2_O_2_) oxidizes Cys_199_, which forms a disulfide bond with neighboring Cys_208_ (19). The oxidized form undergoes a conformational change and contacts four sites on DNA, causing cooperative binding with RNA polymerase to induce gene expression (20, 21). In the reduced form, OxyR contacts 2 different sites on specific DNA sequences to act as a repressor (20). Studies on OxyR homologs in other bacteria have revealed great variation in how OxyR is activated, the genes it regulates, and whether it activates, represses, or both activates and represses genes (22–24). In *E. coli*, approximately 40 genes are in the OxyR regulon, many of which function to reduce oxidative stress in the cell (18). Specifically, genes encoding catalase and alkyl hydroperoxide reductase are under control of OxyR in most Gammaproteobacteria (18). In Alphaproteobacteria, which is the group *Z. mobilis* belongs to, OxyR regulation is more species-specific (18).

Bacterial mechanisms to resist ROS are important because these reactive molecules can damage proteins through oxidation, destabilize the cell membrane through lipid peroxidation, and induce DNA damage and mutations (25). *Z. mobilis* contains a gene encoding an OxyR homolog (*ZMO1733*), but its regulatory role has not been studied. We hypothesize that this homolog regulates the transcriptional response to oxygen-induced stress. In this study, we deleted the gene encoding *oxyR* from the genome of *Z. mobilis* strain ZM4. To characterize the regulon of this transcription factor, we performed RNA-sequencing (RNA-seq) analysis on *ΔoxyR* and wild-type ZM4, grown in oxic and anoxic conditions. We investigated growth of the *oxyR* deletion strain in lignocellulosic hydrolysate and discovered that OxyR is important for growth in this medium. The remainder of the study focuses on uncovering the role of *oxyR* in lignocellulosic hydrolysate fermentation.

## Results

To study the role of OxyR in *Z. mobilis*, we constructed a markerless *oxyR* deletion strain from wild-type ZM4. We also constructed a vector to express the *oxyR* gene for complementation using pRL814, a broad host range vector that bears *gfp* (green fluorescent protein) under a promoter inducible by isopropyl-beta-D-thiogalactoside (IPTG). We replaced *gfp* in pRL814 with *oxyR* to create pRL814-*oxyR*. ZM4 pRL814, *ΔoxyR* pRL814, ZM4 pRL814-*oxyR* (ZM4 *oxyR*), and *ΔoxyR* pRL814-*oxyR* (*ΔoxyR oxyR*) were grown under various conditions to assess the role of *oxyR* in growth. We previously observed leaky expression of *gfp* from this vector in *Z. mobilis*, therefore IPTG was not added in this study (data not shown). In anoxic conditions, deletion of *oxyR* did not impact growth (**Figure 1A**). In oxic conditions, *ΔoxyR* shows a significant growth defect (**Figure 1B**). This is expected, because the canonical role of OxyR is to mitigate the reactive byproducts of aerobic respiration. The complemented strain, *ΔoxyR* pRL814-*oxyR* (labeled Δ*oxyR oxyR* in figures), displays growth similar to wild-type, demonstrating that complementing *oxyR* into the deletion strain background restores growth. From this result, we conclude OxyR is required for robust growth with oxygen in *Z. mobilis*.

**Figure 1.**
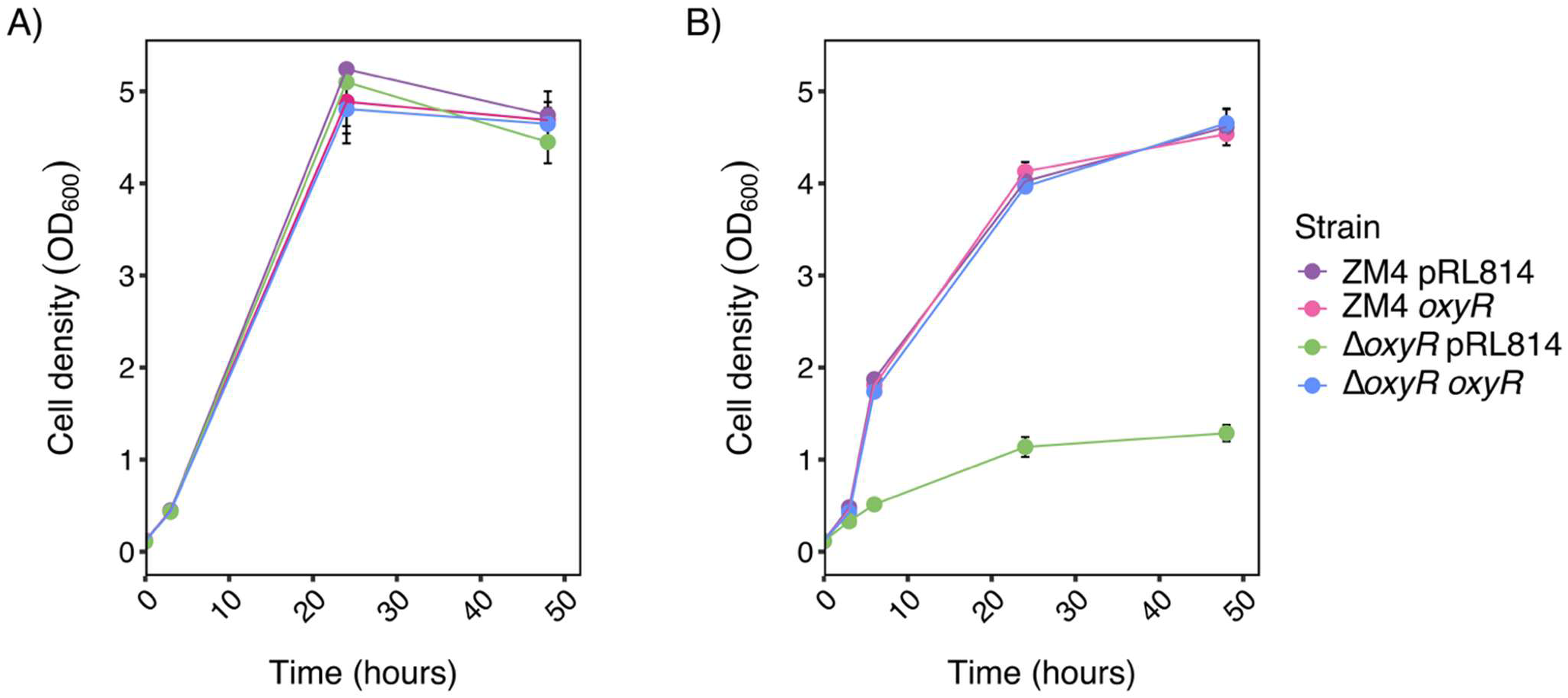
Deletion of *oxyR* inhibits growth under oxic, but not anoxic, conditions. ZM4 pRL814 (purple), ZM4 pRL814-*oxyR* (ZM4 *oxyR*, pink)*, ΔoxyR* pRL814 (green), and *ΔoxyR* pRL814-*oxyR* (*ΔoxyR oxyR*, blue) were grown in anoxic **(A)** and oxic conditions **(B)** for 48 h in ZRMG (rich medium). Points represent the average of 3 biological replicates, and error bars represent standard error.

To further examine the role of OxyR in *Z. mobilis*, we aimed to measure ROS generated in *Z. mobilis*. We used CellROX, an intracellular redox probe, to assess intracellular ROS levels. This dye diffuses across cell membranes and emits fluorescence when oxidized. The CellROX structure is not publicly available, but *in vitro* tests have revealed that it can be oxidized by O^2–^and •OH (26). Strains without the pRL814 plasmid were used as controls in these experiments because the base vector causes GFP production, which could interfere with the fluorescence measurements. Menadione (2-methyl-1,4-naphthoquinone) was used as a positive control because it generates ROS via the formation of semiquinone radicals (27).

Under oxic conditions in rich medium, there was a significant increase in CellROX fluorescence in *ΔoxyR* compared to ZM4 (Tukey-adjusted p = 0.0034) (**Figure 2**). Additionally, *ΔoxyR* pRL814-*oxyR* displayed similar fluorescence to ZM4 and ZM4 pRL814-*oxyR*. This indicates that *Z. mobilis* accumulates intracellular ROS during oxic growth, and that *oxyR* regulation is necessary for effective ROS removal. To assess if *oxyR* is important for ROS resistance in anoxic conditions, the strains were grown anoxically with increasing concentrations of hydrogen peroxide (H_2_O_2_) (**Figure 3**). ZM4 could tolerate up to 1 µM H_2_O_2,_ while the tolerance of *ΔoxyR* was reduced to 0.5 µM H_2_O_2_. ZM4 *oxyR* and *ΔoxyR oxyR* could also tolerate 1 µM but showed a greater lag in growth than ZM4. Growth of these strains at 1 µM H_2_O_2_ was highly variable across replicates, suggesting that this concentration is near a tolerance threshold. Combined, these results indicate that *oxyR* regulation is important for resistance to endogenous ROS generated during aerobic respiration, as well as exogenous ROS during anoxic fermentation.

**Figure 2.**
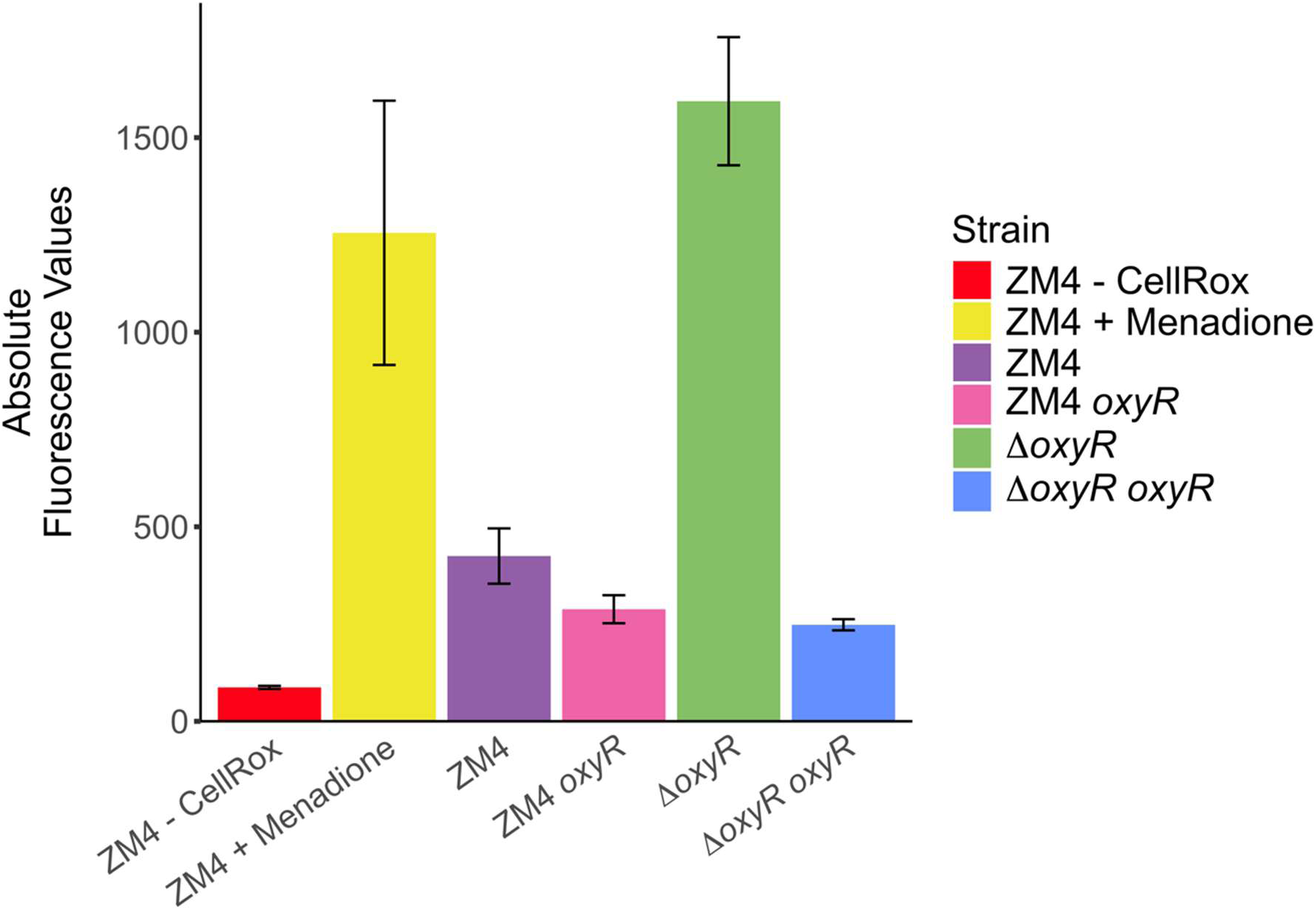
Expression of *oxyR* is essential for removal of intracellular ROS. A CellROX assay was used to quantify oxidant activity after 24 h of oxic growth in ZRMG. Cells were washed in PBS and brought to an OD_600_ of 1. The absolute fluorescence values of ZM4 (purple), ZM4 pRL814-*oxyR* (ZM4 *oxyR*, pink)*, ΔoxyR* (green), and *ΔoxyR* pRL814-*oxyR* (Δ*oxyR oxyR*, blue) were measured at 480 nm excitation, 520 nm emission after incubation with 25 µM CellRox for 30 min. 100 µM menadione was added to ZM4 20 minutes prior to CellROX addition as a positive control for ROS (yellow). No CellRox was added to ZM4 as a negative control (red). Bars represent the average of 3 biological replicates, and error bars represent standard error.

**Figure 3.**
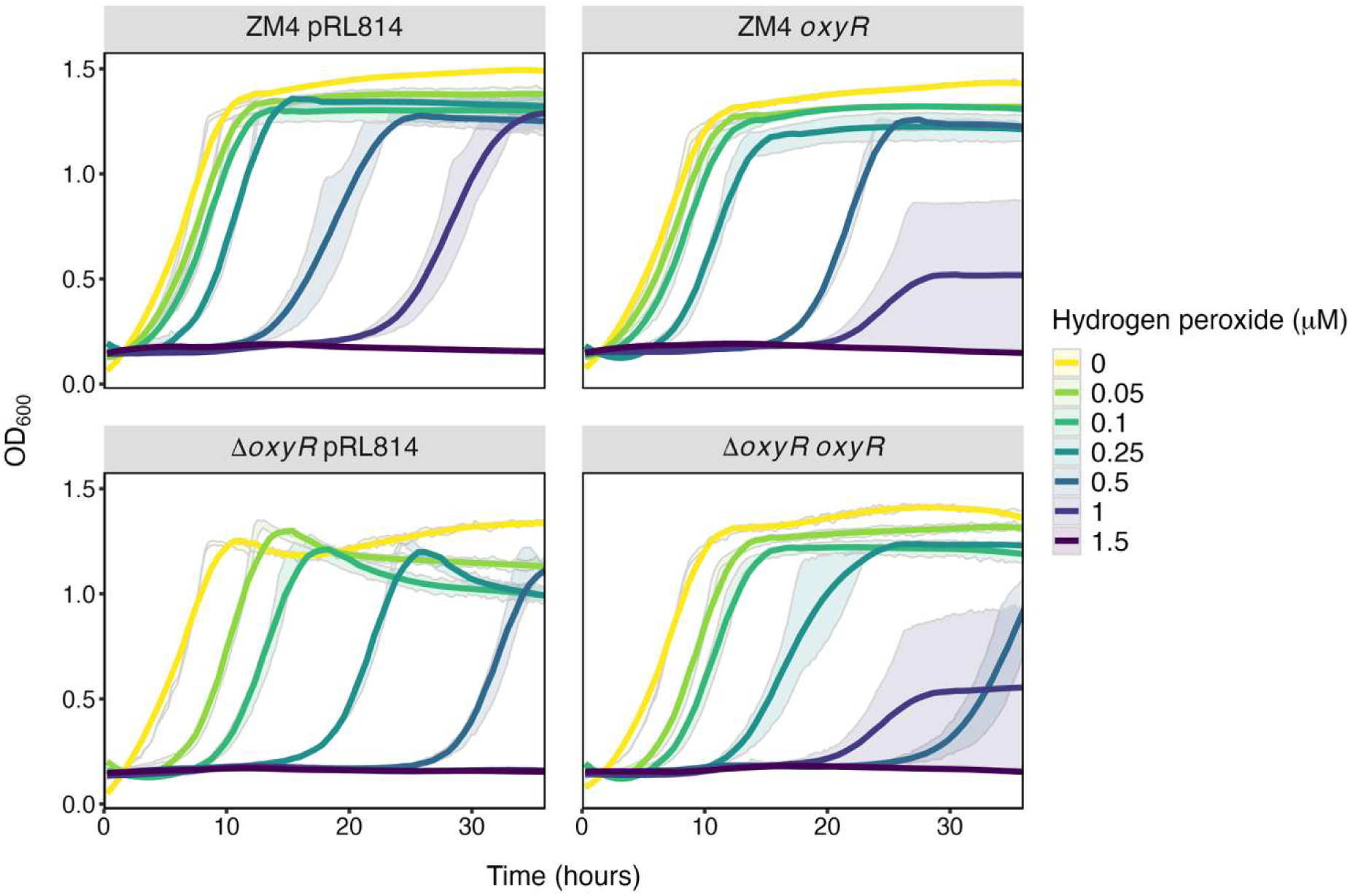
Disruption of *oxyR* decreases resistance to hydrogen peroxide in anoxic conditions. ZM4 pRL814, ZM4 pRL814-*oxyR* (ZM4 *oxyR*)*, ΔoxyR* pRL814, and *ΔoxyR* pRL814-*oxyR* (*ΔoxyR oxyR*) were grown anoxically in ZRMG for 36 hours in concentrations of H_2_O_2_ ranging from 0 – 1.5 µM. Lines represent the average of 3 biological replicates, and standard error values are displayed with transparent ribbon.

To better understand the regulatory role of *oxyR*, we conducted an RNA-seq analysis on ZM4 and *ΔoxyR*, grown in oxic and anoxic conditions (**Figure 4**). Between ZM4 grown in oxic conditions (OX ZM4) and anoxically grown ZM4 (AN ZM4), there were 101 differentially expressed genes. There were 397 differentially expressed genes in OX *ΔoxyR* compared to OX ZM4, representing ∼20% of transcripts. There were also 198 differentially expressed genes in AN *ΔoxyR* compared to AN ZM4. **Figure S1** shows a heatmap of all differentially expressed genes in this dataset. We conclude that deletion of *oxyR* causes global changes to the transcriptome, both in oxic and anoxic conditions. This suggests that OxyR has a broad regulatory role in *Z. mobilis*. Because genes both decrease and increase with *oxyR* deletion, OxyR could be both activating and repressing genes in *Z. mobilis*. Further, we identified several transcriptional regulators that were differentially expressed with *oxyR* deletion (*ZMO0153*, *ZMO0221*, *ZMO1107*, *ZMO1202*, *ZMO1387*, *ZMO1412*, *ZMO1816*), which could alter expression of many downstream genes. It is important to note that these results reflect the indirect regulon of OxyR. An indirect regulon consists of all genes that are impacted when a regulator is deleted, but the regulator may not control the promoters of all directly (28). A direct regulon consists of the genes for which the regulator binds to the promoters to directly affect transcription (28). DNA-binding experiments such as DAP-seq or ChIP-seq would be needed to distinguish between the direct and indirect regulon of OxyR.

**Figure 4.**
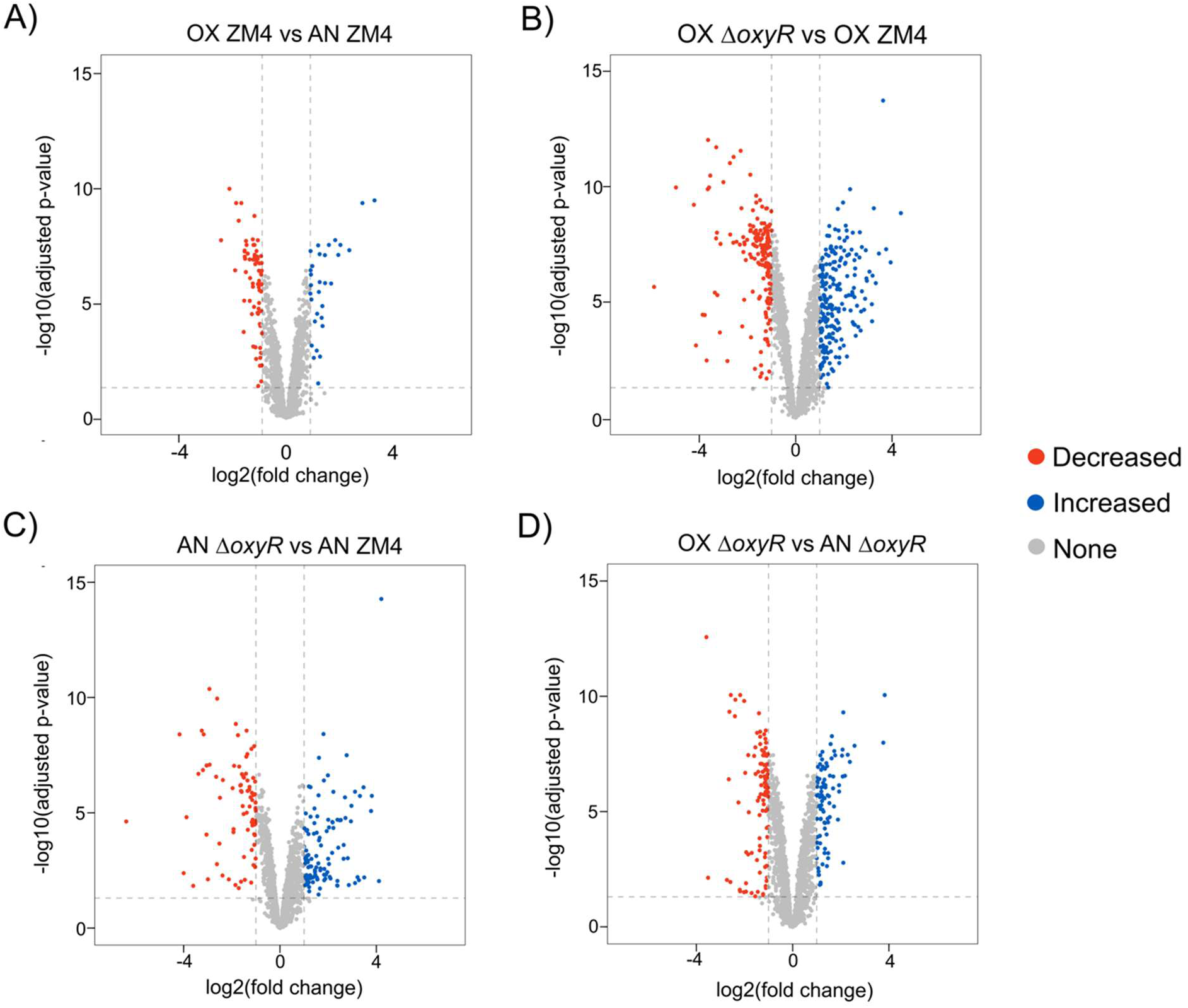
Deletion of *oxyR* causes global transcriptomic reprogramming. Volcano plots show genes that are significantly decreased and increased (> +1 or < -1 log_2_ fold change and False Discovery Rate (FDR) < 0.05) between **A)** oxic ZM4 vs anoxic ZM4, **B)** oxic *ΔoxyR* vs oxic ZM4, **C)** anoxic *ΔoxyR* vs anoxic ZM4, and **D)** oxic *ΔoxyR* vs anoxic *ΔoxyR*.

Once the differentially expressed genes (DEGs) were visualized at the global scale, the genes that are likely regulated by OxyR in different conditions were identified based on log_2_ fold change between groups. **Figure 5A** shows significant DEGs (> +1 or < -1 log_2_ fold change and False Discovery Rate (Benjamini-Hochberg adjusted p-value, FDR) < 0.05) that were the most upregulated in Δ*oxyR* under both oxic and anoxic conditions. These are genes that OxyR likely represses under all conditions. Interestingly, prophage genes that are present on one of the *Z. mobilis* plasmids made up most of these genes. There are also two genes in the Cell Wall category (*ZMO0932* and *ZMOp36×001*) that encode enzymes that break down peptidoglycan. **Figure 5B** shows significant DEGs that were the most downregulated in Δ*oxyR* under both conditions. These genes are likely consistently activated by OxyR. Many flagellar biosynthesis genes (*ZMO0607*-*ZMO0614*, *ZMO0632*, and *ZMO0634*) belong to this category, indicating that OxyR could play a role in promoting motility. **Figure 5C** shows genes with the largest differential response to *oxyR* deletion between oxic and anoxic conditions. Specifically, genes that simultaneously were downregulated in *ΔoxyR* compared to ZM4 in oxic conditions and upregulated in *ΔoxyR* compared to ZM4 in anoxic conditions. These are genes that are presumed to be activated by OxyR during oxic growth and repressed by OxyR in anoxic growth. There are several genes encoding redox enzymes in this category, such as ferredoxin-NADP reductase, which catalyzes the transfer of electrons between NADPH and ferredoxin. A key gene regulated by OxyR in other organisms, *ahpC* (*ZMO1732*), is present in this category. This gene encodes the peroxiredoxin alkyl hydroperoxide reductase AhpC, which reduces organic peroxides to their respective alcohols and plays a crucial role in the bacterial response to ROS. This gene had a -1.89 log_2_ (∼3.5x) fold change in oxic Δ*oxyR* vs oxic ZM4. Additionally, *ahpC* had a 2.62 log_2_ (∼6.2x) fold change in oxic ZM4 vs anoxic ZM4. Also in this category is *ZMO1060,* encoding superoxide dismutase, another key enzyme canonically regulated by OxyR. Superoxide dismutase plays a key role during oxidative stress through the dismutation of superoxide radical (O_2_ ^-^) into less toxic O and H O. ETC components NADH dehydrogenase and lactate dehydrogenase are also in this group and may increase expression during oxidative stress to reduce oxygen and limit the formation of ROS. The largest category of genes seemingly activated by OxyR during oxic growth is sulfur metabolism. Genes *cysN* (*ZMO0004*), *cysD* (*ZMO0005*), and *cysG* (*ZMO0006*) belong to the Cysteine biosynthesis *cysCND* operon. Together, *cysI* (*ZMO0008*) and *cysJ* (*ZMO0009*) encode the two subunits that make up sulfite reductase, which reduces sulfite to hydrogen sulfide in the assimilatory sulfate reduction pathway (29). This suggests that OxyR plays an important role in regulating sulfur metabolism in the presence of oxygen.

**Figure 5.**
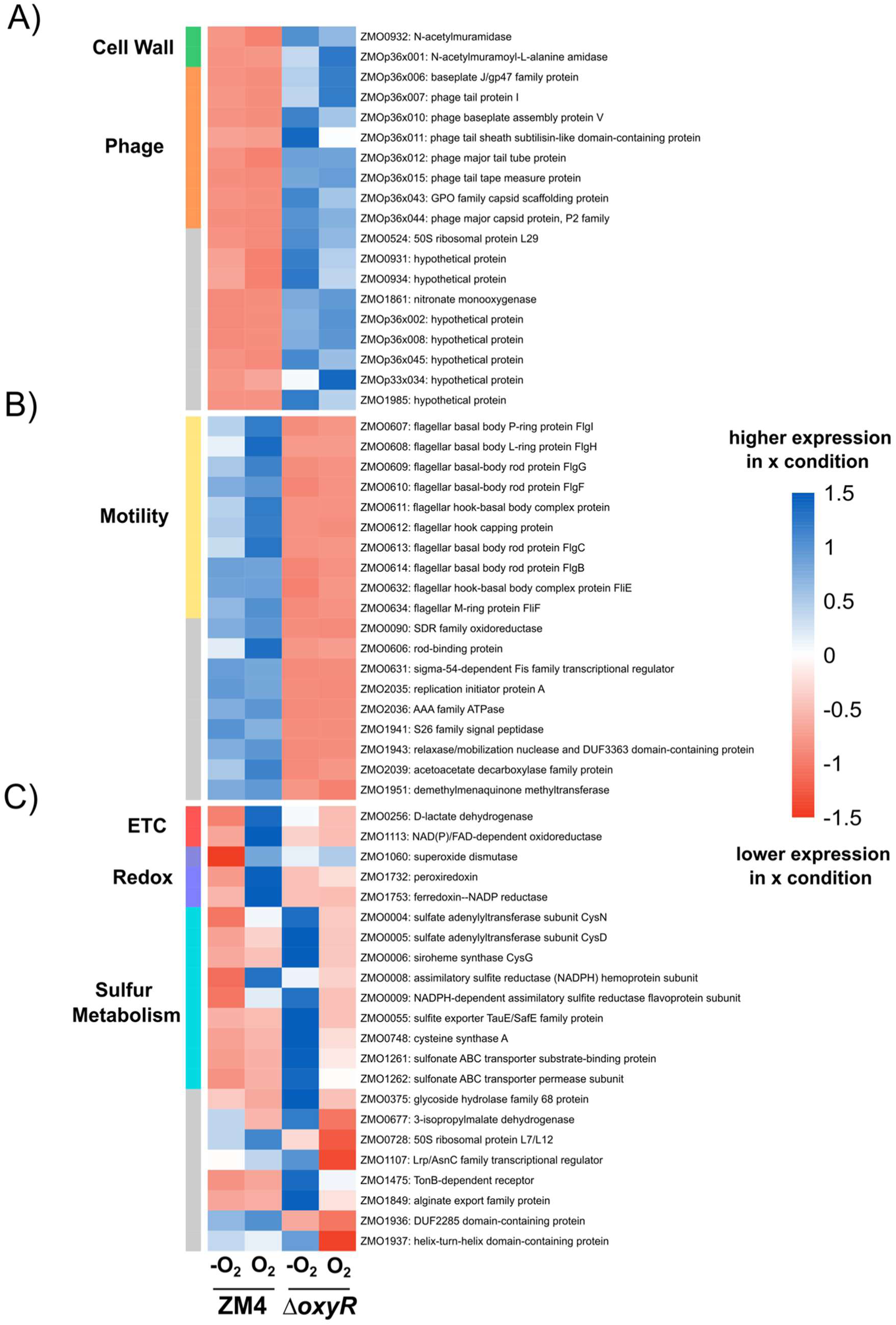
Differential regulation of genes in Δ*oxyR*. Heatmaps show the row-centered z-scores of the Fragments Per Kilobase Million (FPKM) value for each gene in the four conditions. **A)** Significant differentially expressed genes (DEGs) that were upregulated in *ΔoxyR* across both conditions. DEGs were defined as genes that had significant differential expression (> +1 or < -1 log_2_ fold change and False Discovery Rate (Benjamini-Hochberg adjusted p-value, FDR) < 0.05) in at least one contrast between conditions. Genes were selected that showed greater than 2 log_2_ fold change in oxic *ΔoxyR* vs oxic ZM4, as well as in anoxic *ΔoxyR* vs anoxic ZM4. **B)** Significant DEGs that were downregulated in *ΔoxyR* across both conditions. Genes were selected that showed less than -2 log_2_ fold change in oxic *ΔoxyR* vs oxic ZM4, as well as in anoxic *ΔoxyR* vs anoxic ZM4. **C)** Genes with the largest differential response to *oxyR* deletion between anoxic and oxic conditions. For each significant DEG, the difference between the log_2_ fold change in oxic *ΔoxyR* vs oxic ZM4 was subtracted from the log_2_ fold change in anoxic *ΔoxyR* vs anoxic ZM4. Genes that had a value greater than 1.5 in this metric are shown. In each heatmap, genes that had similar functions were grouped together. The gray bars indicate the genes that do not belong to a shared category. FPKM values were calculated as the average across three or four biological replicates.

To assess whether *oxyR* regulation is important in a biofuel production context, the ZM4 and *oxyR* mutant strains were grown in 7% Ammonia-Fiber Expansion (AFEX) switchgrass hydrolysate (**Figure 6**). This is hydrolysate that was produced from switchgrass biomass using the AFEX method of pretreatment, and contains 7% glucan content (30, 31). The AFEX pretreatment method consists of adding liquid ammonia to harvested switchgrass at high temperature and pressure (30, 31). AFEX pretreatment causes the decrystallization of cellulose, partial depolymerization of hemicellulose, cleavage of lignin-carbohydrate complex linkages, and increases accessible surface area for hydrolytic enzymes (32). Enzymes such as cellulases are added to the plant biomass to depolymerize the plant sugars in a process called hydrolysis (31). This process generates the liquid hydrolysate containing high concentrations of glucose and xylose, along with many compounds inhibitory for growth. AFEX is a preferred method of pretreatment because it generates lower concentrations of inhibitory compounds than other methods (33).

**Figure 6.**
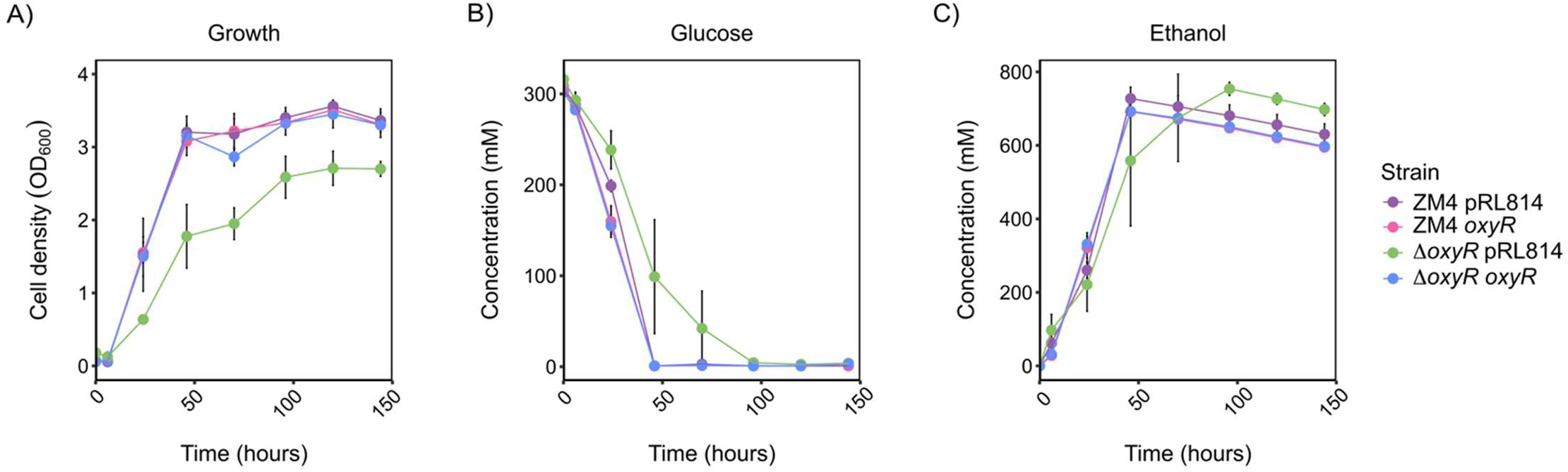
Growth of *ΔoxyR* in 7% AFEX switchgrass hydrolysate. **A)** Anoxic growth of ZM4 pRL814 (purple), ZM4 pRL814-*oxyR* (ZM4 *oxyR*, pink)*, ΔoxyR* pRL814 (green), and *ΔoxyR* pRL814-*oxyR* (*ΔoxyR oxyR*, blue) after 144 hours in 7% AFEX switchgrass hydrolysate. **B-C)** At each time point in the growth curve, samples were taken for HPLC analysis, and the concentrations of glucose and ethanol were determined. Points represent the average of 3 biological replicates, and error bars represent standard error.

In hydrolysate, *ΔoxyR* pRL814 has a growth defect compared to ZM4 pRL814, although not as pronounced as in oxic conditions (**Figure 6A**). Growth was rescued in the complementation strain. *ΔoxyR* pRL814 also took longer to consume glucose and produce ethanol in hydrolysate (**Figure 6B and C**). It is interesting that despite the lower OD_600_ observed in *ΔoxyR* pRL814, this strain was still able to consume all glucose and convert it to ethanol. Wild-type ZM4 produces the maximal amount of ethanol (727 ± 31 mM) after 48 hours of growth in hydrolysate. It takes 96 hours for the *ΔoxyR* pRL814 strain to produce a comparable ethanol concentration (754 ± 18 mM). These results indicate that although OxyR regulation is canonically important for oxic growth, it is also important for efficient, anoxic biofuel production from hydrolysate in *Z. mobilis*.

To further investigate the effect of hydrolysate components on growth of the *ΔoxyR,* strains ZM4 and *ΔoxyR* were grown anoxically in diluted and undiluted hydrolysate in a 96-well plate (**Figure 7**). In undiluted hydrolysate, the average lag time to reach an OD_600_ of 0.1 above the starting OD_600_ was 17.7 ± 0.14 hours for ZM4 and 20.7 ± 2.29 hours for *ΔoxyR*. With the dilution of hydrolysate by half, the average lag times decreased to 3.6 ± 0.54 (Tukey-adjusted *p* = 0.00013) for ZM4 and 2.6 ± 0.08 hours (Tukey-adjusted *p* = 2.18 x10^-5^) for *ΔoxyR*. While the dilution decreased the lag time of *ΔoxyR* to be on par with diluted ZM4, *ΔoxyR* continued to have a lower maximum OD_600_ than ZM4 in both conditions. However, the difference in maximum OD_600_ was only significant between ZM4 and *ΔoxyR* in undiluted hydrolysate (mean difference = 0.49, Tukey-adjusted *p* = 0.0011), not 50% diluted hydrolysate (mean difference = 0.25, Tukey-adjusted *p* = 0.15). These results confirm that components of hydrolysate are more inhibitory to *ΔoxyR* than ZM4, and when these components are diluted, the inhibitory effect is lessened.

**Figure 7.**
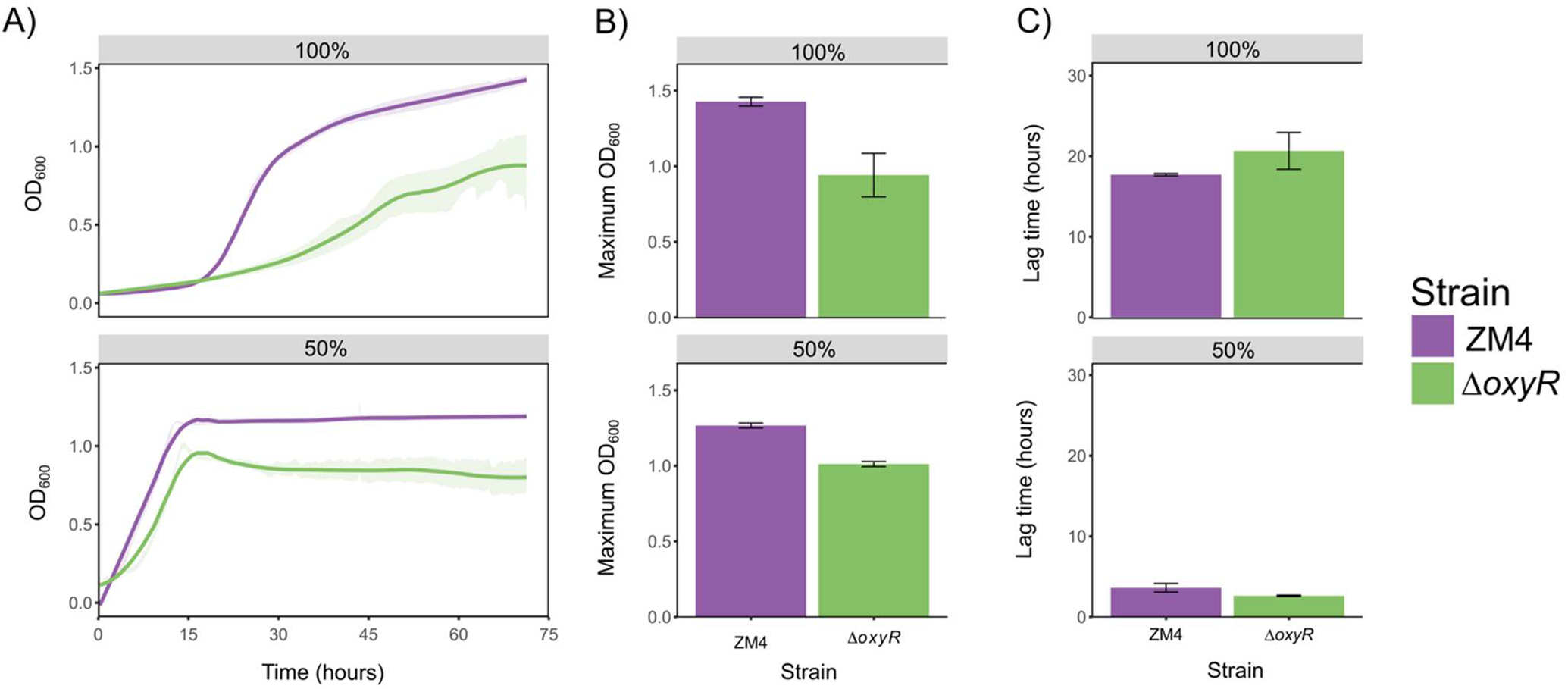
Dilution of 7% AFEX switchgrass hydrolysate accelerates growth of *ΔoxyR*. ZM4 (purple) and *ΔoxyR* (green) were grown anoxically in AFEX hydrolysate in a 96-well plate for 72 hours in undiluted hydrolysate (100%) and hydrolysate diluted by half with sterile water (50%). **A)** Growth of the strains in undiluted and diluted hydrolysate over time. Lines represent the average of 3 biological replicates, and standard error values are displayed with transparent ribbon. **B)** Maximum OD_600_ the strains reached during the time course. Bars represent the average of 3 biological replicates, and error bars represent the standard error. **C)** Average lag time of each strain in undiluted and diluted hydrolysate. Lag time was calculated as the time to reach 0.1 OD_600_ above the starting OD_600_. Bars represent the average of 3 biological replicates, and error bars represent the standard error.

Lignocellulosic hydrolysate contains diverse growth inhibitors that are formed during pretreatment, such as organic acids (acetic acid, formic acid), aldehydes (furfural, 5-hydroxymethylfurfural [HMF]), and phenolic cinnamic acids (ferulic acid, coumaric acid) (2). Many of these compounds are also partially converted to derivatives such as amides and alcohols (2). We hypothesized that either the synergistic effect of these inhibitors or specific inhibitors could cause oxidative stress in *Z. mobilis*, which could explain why deletion of *oxyR* hinders growth. Many of these inhibitory compounds are known to cause oxidative stress in bacteria in oxic conditions, but it is unknown whether they have the same effect in anoxic conditions. We first grew *ΔoxyR* pRL814 in standard lab media with specific hydrolysate inhibitors that were identified to cause a growth defect in *oxyR* mutants in a chemical genomics CRISPRi library screen when high concentrations were used (34). No significant growth inhibition was observed in *ΔoxyR* pRL814 when grown anoxically in concentrations similar to those found in hydrolysate (**Figure S2**). This result indicates that individual inhibitory chemicals are not responsible for the observed growth defect in hydrolysate. To assess whether a synergistic effect of many compounds could be responsible, we used a synthetic hydrolysate that mimics AFEX switchgrass hydrolysate, SynHv4.1. This hydrolysate contains the same concentrations of glucose, xylose, and many lignocellulose-derived growth inhibitors that are present in AFEX hydrolysate and was prepared as described in Barten et al. (35). A full list of these compounds and their concentrations can be found in Supplementary Table 1. The strains were anoxically grown in SynHv4.1 for 72 hours. Unlike in the AFEX hydrolysate, there was no growth defect of the *ΔoxyR* strain in SynHv4.1 (**Figure 8**). This indicates that factors other than the known inhibitors in AFEX hydrolysate cause growth inhibition when *oxyR* is disrupted. Additionally, all strains grew faster in SynHv4.1 than in AFEX hydrolysate. This suggests that there are yet unknown components in the AFEX hydrolysate that impact *Z. mobilis* growth.

**Figure 8.**
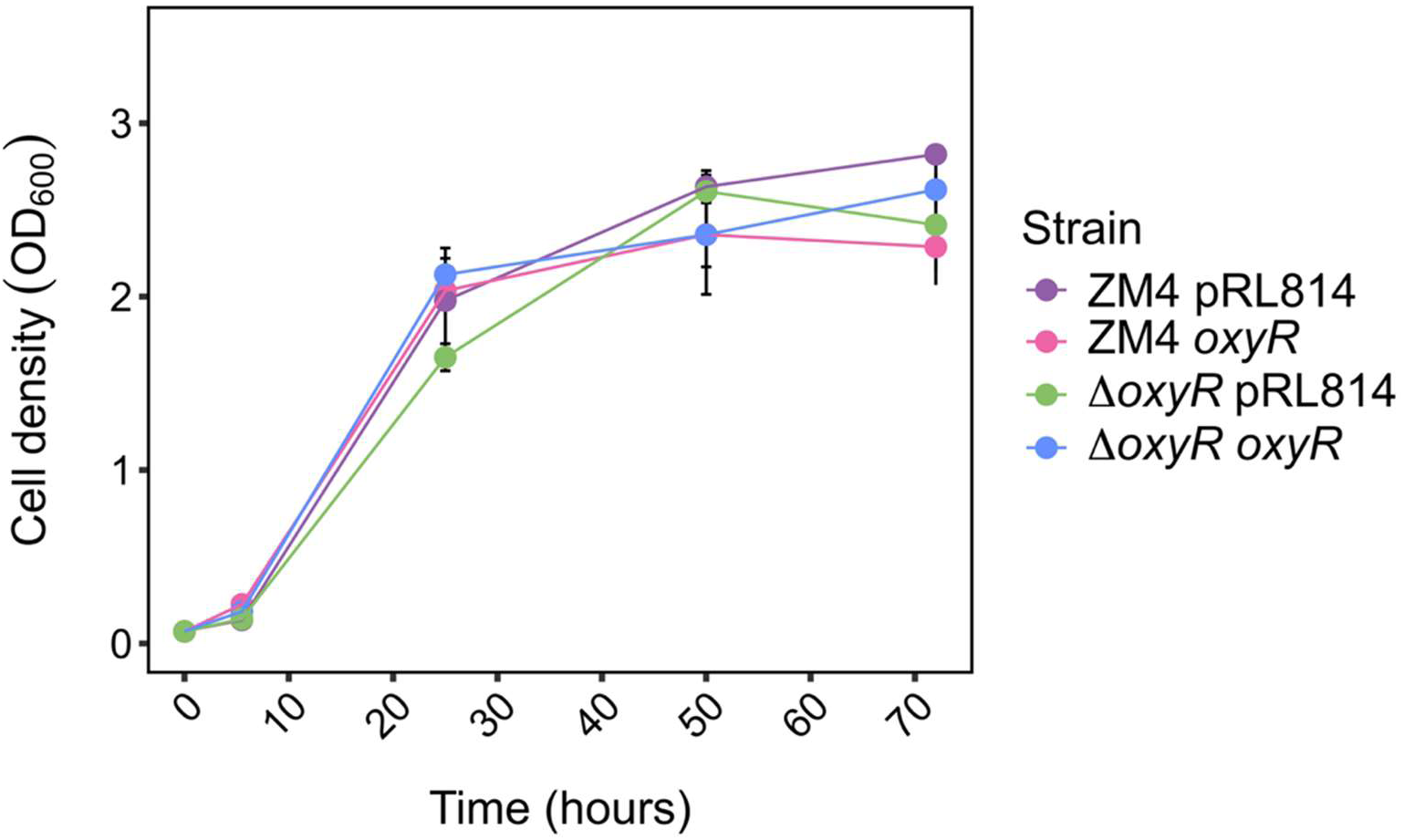
Growth of *ΔoxyR* in synthetic hydrolysate SynHv4.1 shows no growth defect. ZM4 pRL814 (purple), ZM4 pRL814-*oxyR* (ZM4 *oxyR*, pink)*, ΔoxyR* pRL814 (green), and *ΔoxyR* pRL814-*oxyR* (Δ*oxyR oxyR*, blue) were grown anoxically for 72 hours in synthetic hydrolysate SynHv4.1. Points represent the average of 3 biological replicates, and error bars represent standard error.

A peroxide assay was used to quantify the concentration of peroxides present in both hydrolysates. This assay relies on the FOX method (ferrous oxidation in xylenol orange), in which peroxides or other strong oxidants present in a sample oxidize ferrous iron (Fe^2+)^ to ferric iron (Fe^3+^) (36). Fe^3+^ binds to xylenol orange, forming a purple complex that can be measured at 560 nm absorbance. This test cannot differentiate between the exact compounds oxidizing Fe^2+^ (H_2_O_2_, organic peroxides, or other strong oxidants) in a sample, however we are reporting these results as the concentration of peroxides present in the medium, as this is what the assay is intended to measure. The concentrations were calculated based on an external standard curve generated with H_2_O_2_. Peroxide concentrations were measured in AFEX hydrolysate, SynHv4.1, and ZRMG, using samples of these media that had been oxic or anoxic for over two months. Oxic AFEX and SynHv4.1 hydrolysates had average peroxide levels of 49 µM and 16 µM, respectively. Anoxic AFEX and SynHv4.1 hydrolysates had no detectable peroxides (**Figure 9**). ZRMG had no detectable peroxide levels in either oxic or anoxic conditions (**Figure 9**). We then placed the oxic samples in the anaerobic chamber, and the anoxic samples in the oxic atmosphere (benchtop). The peroxide levels were measured each day following this switch. Oxic SynHv4.1 peroxide levels dropped to zero within two days in the anaerobic chamber, while it took eight days for AFEX hydrolysate to do the same. After eight days, the anoxic hydrolysates reached similar starting concentrations measured in the oxic hydrolysates (**Figure 9**).

**Figure 9.**
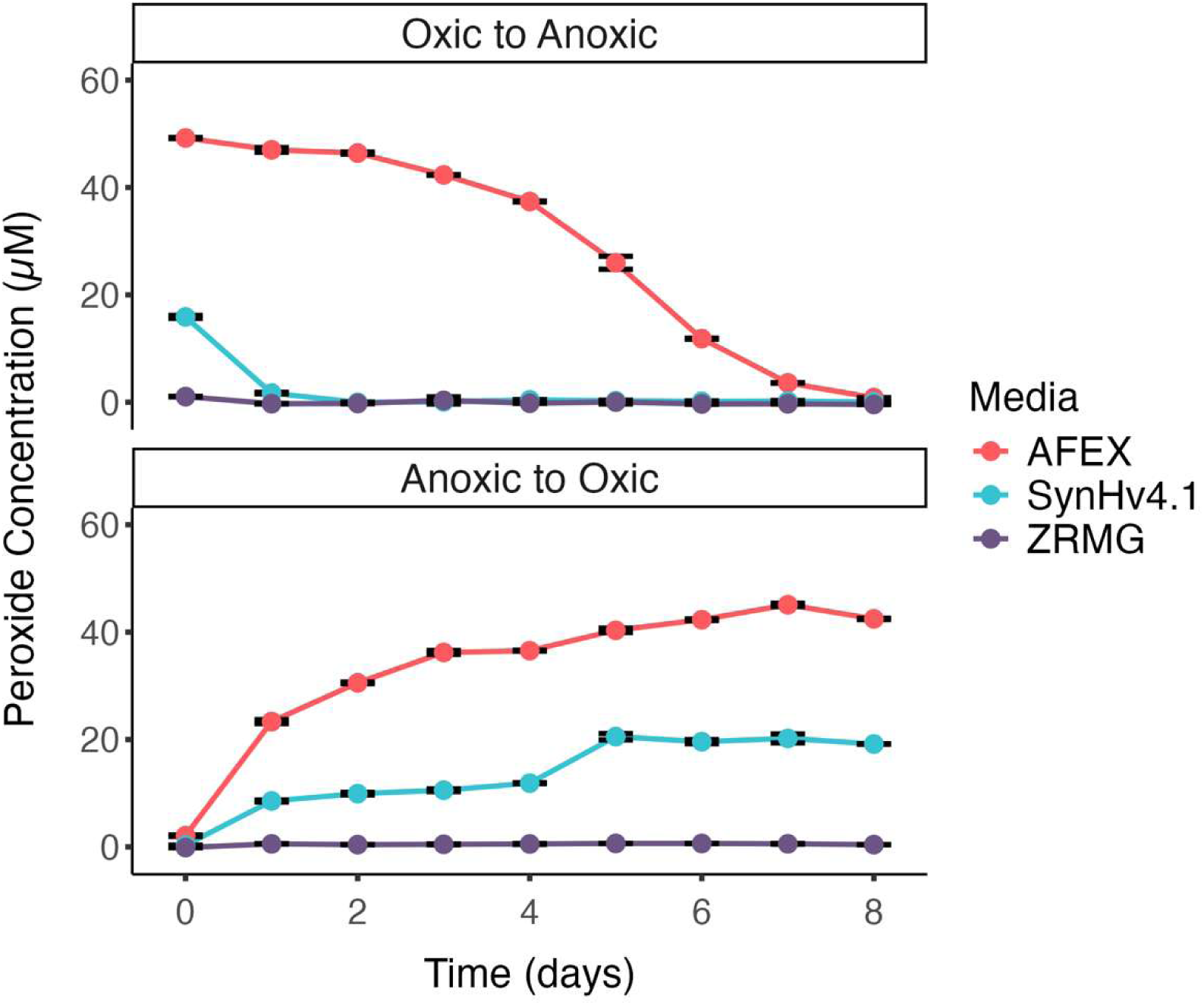
AFEX hydrolysate contains significant peroxide levels that persist over time. The Pierce Peroxide Assay was used to measure peroxide levels in AFEX and SynHv4.1 synthetic hydrolysate. ZRMG was used as a control. Media started out as either anoxic or oxic and was switched to the opposite oxygen condition on Day 0. Absorbance was measured at 595 nm every 24 hours. Points represent the average of 3 technical replicates, and error bars represent standard error.

We next explored whether the peroxides present in AFEX hydrolysate limit growth. ZM4 and Δ*oxyR* were grown anoxically in AFEX hydrolysate for 72 hours in a 96-well plate (**Figure 10**). The addition of exogenous catalase from bovine liver reduced the lag time of both strains. *ΔoxyR* had a significant decrease in lag time at catalase 0.5 U/mL compared to 0 U/mL (mean difference = 8.44 h, Tukey-adjusted *p* = 0.00027). Although not as pronounced, ZM4 also had a significant decrease in lag time at catalase 0.5 U/mL compared to 0 U/mL (mean difference = 4.84 h, Tukey-adjusted *p* = 0.024). Even with the addition of catalase, ZM4 still had faster and higher overall growth than Δ*oxyR*, likely because Δ*oxyR* regulates a complex response that includes more than catalase alone. Additionally, catalase converts H_2_O_2_ into water and O_2_, which would still cause inhibitory effects on Δ*oxyR*. Interestingly, both ZM4 and Δ*oxyR* reached higher maximum OD_600_ by the end of the growth curve without the addition of catalase, despite their longer lag times (**Figure 10**). Overall, these results suggest that peroxides contribute to Δ*oxyR* inhibition in hydrolysate, as well as the slow growth of wild-type *Z. mobilis* in hydrolysate.

**Figure 10.**
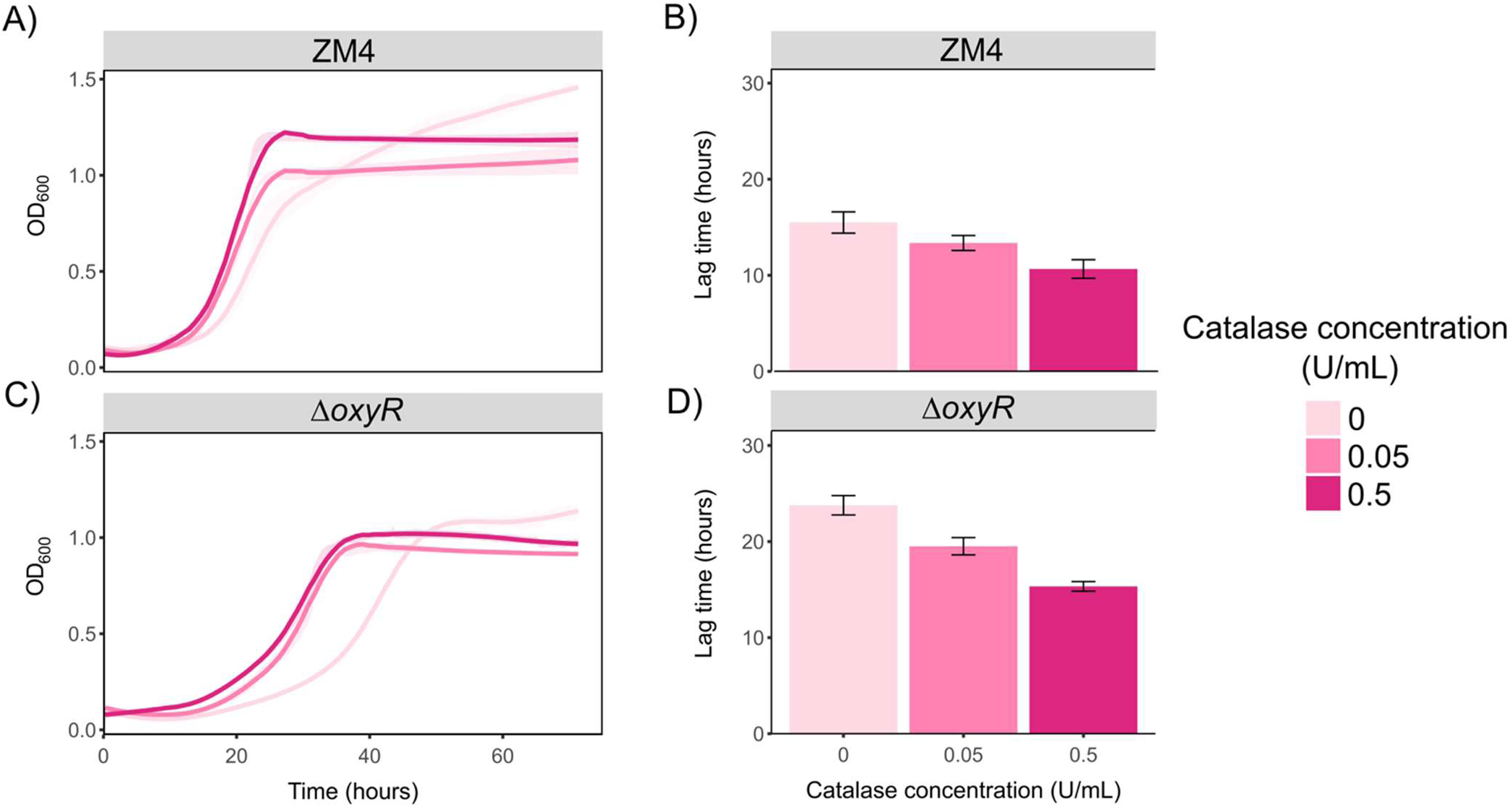
Catalase reduces lag time in hydrolysate for *ΔoxyR* and ZM4. Strains were grown anoxically in AFEX hydrolysate in a 96-well plate for 72 hours, supplemented with 0, 0.05, or 0.5 Units/mL of catalase enzyme. **A and C)** Growth of the strains over time in different catalase concentrations. Lines represent the average of 3 biological replicates, and standard error values are displayed with transparent ribbon. **B and D)** Average lag time of each strain in each catalase concentration. Lag time was calculated as the time to reach 0.1 OD_600_ above the starting OD_600_. Bars represent the average of 3 biological replicates, and error bars represent the standard error.

## Discussion

This study aimed to elucidate the role of OxyR in *Z. mobilis*, a transcription factor that regulates the response to oxidative stress. Hydrolysate causes an oxidative stress response in other organisms, such as *Saccharomyces cerevisiae*, suggesting that improving the oxidative stress response could be a general method for enhancing lignocellulosic biofuel production (37). Indeed, a previous study identified a single nucleotide polymorphism (SNP) in *oxyR* in a plasmid-free *Z. mobilis* strain with enhanced biofuel production characteristics (38). This strain showed improved conversion of glucose to ethanol in xylose mother liquor compared with the wild-type ZM4 strain (38). The *oxyR* mutation was responsible for lower ROS levels measured in the mutant strain in addition to its better performance consuming xylose mother liquor (38). In a subsequent study, the authors show that the *oxyR* mutation (OxyR^S267P^) decreased the steric hindrance for disulfide bond formation, making it more prone to transition to the oxidized state (39). The mutated OxyR was used to develop a biosensor for oxidative stress induced by inhibitors in toxic xylose mother liquor (39). In another study from the same group, a different SNP in *oxyR* was found in an acid-tolerant strain of *Z. mobilis* (40). These findings suggest that OxyR could be important for resistance to different types of stress in *Z. mobilis*, including growth in hydrolysate, making OxyR relevant for industrial use of this organism. To carefully assess the role of OxyR in *Z. mobilis*, we constructed an *oxyR* deletion strain and investigated the effects of this deletion. We found that OxyR in *Z. mobilis* has canonical roles, such as enhancing oxic growth and resistance to H_2_O_2_. Interestingly, *ΔoxyR* pRL814-*oxyR* and ZM4 pRL814-*oxyR* had decreased growth in the higher H_2_O_2_ concentrations compared to ZM4 pRL814. This suggests that changing the natural levels of OxyR to any degree can impact ROS tolerance in *Z. mobilis*.

We performed an RNA-seq analysis of the *oxyR* deletion strain to determine the broad OxyR regulon (direct and indirect). The genes that OxyR likely activates in response to oxidation have functions in redox activity, the ETC, and sulfur metabolism. One of the genes found in the redox category is *ahpC,* encoding peroxiredoxin. Other oxidative stress response enzymes regulated by OxyR in *E. coli*, such as glutaredoxin and catalase, did not have significant differential expression in *Z. mobilis* Δ*oxyR*. Thus, AhpC may play the main role in reducing ROS in *Z. mobilis*. In *E. coli*, AhpC is the primary scavenger of H_2_O_2_ (41). The gene encoding AhpC is also adjacent to *oxyR* in the *Z. mobilis* genome, further suggesting its importance in the OxyR regulon. Similarly, Wang et al. found that the *Z. mobilis* strain containing the OxyR^S267P^ mutation had greater gene expression of *ahpC* compared to the reverted strain containing the wild-type *oxyR* gene (39). We over-expressed *ahpC* in Δ*oxyR* but did not observe a significant increase in oxic growth compared to Δ*oxyR* pRL814 (**Figure S3A**). Additionally, during oxic growth, Δ*oxyR* pRL814-*ahpC* had similar CellROX fluorescence to Δ*oxyR* pRL814 (**Figure S3B**). Perhaps the expression of *ahpC* alone is not sufficient to restore ROS levels to wild-type, because other genes normally activated by OxyR are still lacking in the deletion strain. AhpC reduces organic peroxides using a Cysteine residue in its active site, which then forms an intramolecular disulfide bond with a neighboring Cysteine (42). AhpF, alkyl hydroperoxide reductase subunit F, reduces the disulfide bond in AhpC using NAD(P)H so that AhpC can reduce other peroxide substrates (42). In bacteria that lack AhpF, the thioredoxin system of thioredoxin reductase (TrxR) and thioredoxin (Trx) mediates the recycling of AhpC (42). There is no gene annotated in *Z. mobilis* as encoding AhpF, but both thioredoxin and thioredoxin reductase genes are present in the genome, so it is likely that the thioredoxin system recycles AhpC using NAD(P)H.

ETC components NADH dehydrogenase (*ndh*) and lactate dehydrogenase (*ldh*) appear to be activated by OxyR aerobically, and they were both previously found to be upregulated in *Z. mobilis* in response to oxygen (14). Upregulation of ETC dehydrogenases could be a strategy *Z. mobilis* uses to mitigate ROS. The *Z. mobilis* ETC is necessary for its oxic growth because it reduces O_2_ in the cell (16). By increasing expression of components necessary for aerobic respiration during oxidative stress, *Z. mobilis* may increase the rate of oxygen consumption, potentially limiting the amount of ROS formed through electron transfer to O_2_. It is important to note that the ETC can also be a source of oxidative stress in bacteria through the leakage of single electrons that generate H_2_O_2_ and O_2_ ^-^ (43).

Genes related to assimilatory sulfate reduction were also downregulated in oxic Δ*oxyR*. This pathway produces hydrogen sulfide (H_2_S), which acts as a protectant against oxidative stress in bacteria (44). In *E. coli*, H_2_S combats oxidative stress through sequestration of Fe^2+^ to limit the damaging Fenton reaction, in which Fe^2+^ reacts with H_2_O_2_ to form •OH (45). In *E. coli*, H_2_S also enhances the activity of antioxidant enzymes catalase and superoxide dismutase (45). Cysteine biosynthesis genes also appear to be activated by OxyR in oxygen and repressed in anoxic conditions. Cysteine plays a pivotal role in the bacterial ROS response. Cysteine is one of three amino acids that make up the antioxidant Glutathione. Glutathione protects bacteria from oxidative stress by reducing disulfide bonds that have formed in proteins from oxidative damage. Another role of Glutathione is glutathionylation, which is the formation of disulfide bonds between glutathione and protein Cysteine residues that occurs mainly during oxidative stress (46, 47). This is thought to protect proteins from overoxidation of Cysteine residues to sulfinic acid or sulfonic acid, which could result in irreversible inactivation (46). In *E. coli*, the disulfide bond of activated OxyR is re-reduced by glutaredoxin via glutathione (19). Additionally, cysteine supplementation increases *Z. mobilis* growth and decreases ROS accumulation during growth in the presence of furfural, which is an inhibitory compound found in hydrolysate (46, 47). In all, many genes that appear to be activated by OxyR in oxygen have roles in mitigating formation of ROS and regulating intracellular redox balance.

The RNA-seq analysis also revealed genes that appear to be regulated by OxyR regardless of its oxidation status. Many flagellar genes were downregulated in *ΔoxyR*, suggesting that OxyR normally activates (or de-represses) the expression of these genes in both conditions. OxyR was similarly found to positively regulate flagellin and pili genes in *Acidovorax citrulli* (48). OxyR also influences motility via biofilm formation in several pathogenic bacterial species (49–51). Further work is needed to elucidate the role of OxyR in regulating motility in *Z. mobilis*. Many prophage genes were upregulated in *ΔoxyR*, which indicates constitutive repression of these genes by OxyR. OxyR represses both mu and lambda prophage genes in *E. coli*, due to OxyR binding sites near the promoters of both prophage (52, 53). It was found that the active OxyR protein improves binding of a lambda phage repressor to its operator, repressing transcription of the lambda prophage (53). In the mu *mom* operon, it was observed that OxyR binds to the unmethylated promoter and hinders binding of RNA Polymerase (52). Similar mechanisms may be occurring in *Z. mobilis,* and future research should investigate them.

We next discovered that OxyR plays an important role in *Z. mobilis* biofuel fermentation. We found that deletion of *oxyR* is inhibitory during growth in hydrolysate. Our results show that despite synthetic hydrolysate containing the same known chemical components as AFEX hydrolysate, it does not produce a growth defect in the *oxyR* deletion strain. This aligns with our finding that AFEX hydrolysate contains around 50 µM of peroxide after exposure to oxygen, while synthetic hydrolysate contains less than half of that concentration. We propose that peroxide generates intracellular ROS in *Z. mobilis* during hydrolysate growth, which is why the deletion of OxyR is more deleterious in AFEX hydrolysate than SynHv4.1. Supporting this hypothesis, we found that the addition of exogenous catalase to hydrolysate improved the growth of *ΔoxyR*, significantly reducing its lag time. Interestingly, catalase also reduced the lag time of ZM4. Our RNA-Seq analysis found that the *Z. mobilis* catalase gene, *katE* (*ZMO918*) was not significantly differently expressed in the *oxyR* deletion strain, suggesting that catalase is not regulated by OxyR in *Z. mobilis*. These results suggest that the wild-type *Z. mobilis* oxidative stress response in hydrolysate is inadequate to detoxify the peroxides present. This provides opportunities to improve *Z. mobilis* performance in hydrolysate through the addition of catalase or genetic engineering to improve its enzymatic removal of ROS.

Because identification of peroxides in hydrolysate may be useful for strain engineering, it is important to determine whether the peroxide levels measured correspond to H_2_O_2_ or organic peroxides. Addition of catalase to AFEX hydrolysate did not reduce measured peroxide concentrations after 1 h treatment (**Figure S4**). This result indicates that organic peroxides, not H_2_O_2_, are the peroxide species present in hydrolysate. Because catalase only degrades H_2_O_2_ and it improved growth, we can also assume that H_2_O_2_ is generated in *Z. mobilis* when it is exposed to the organic peroxides present. This could be due to the activity of superoxide dismutase, which generates H_2_O_2_. These varying ROS forms must be detoxified by different pathways, therefore, additional characterization of the oxidative stresses found in hydrolysate and those generated in *Z. mobilis* will be helpful.

It remains unknown how peroxides are formed in hydrolysate. Because hydrolysate is produced from plant matter and undergoes the AFEX pretreatment process, it is feasible that processing causes radical compounds and peroxides to form. Polyphenolic compounds, which are present in both the synthetic and real hydrolysate, may act as either antioxidants or pro-oxidants that either reduce or generate ROS (54, 55). It is possible that during the pretreatment process, these polyphenols could form phenoxy radicals. One study found that alkaline conditions caused an increase in radical anions in lignin extracts, and that this reaction proceeded in both anoxic and oxic media (56). The authors concluded that *o*-semiquinone radicals are likely formed in alkaline medium via decomposition of quinone complexes with hydroquinones (quinhydrones) present in lignin (56). A similar mechanism could occur during AFEX pretreatment, in which ammonia penetrates the biomass and reacts with water to form ammonium hydroxide (31). The generation of radical quinone or polyphenolic species by hydroxide ions during pretreatment could be responsible for peroxide formation. This would explain why *ΔoxyR* did not have a growth defect in SynHv4.1, which does not undergo pretreatment.

Tannis are another potential source of the reactive chemical species in hydrolysate. Tannins are plant phenolic compounds with sizes ranging from 500 to 3,000 Da (57). They vary widely in structure. Hydrolysable tannins are composed of a central carbohydrate molecule with ellagic acid or gallic acid groups attached, while condensed tannins are composed of repeating flavonoid units linked through carbon-carbon bonds (57). A study investigating the effect of Black wattle tannin extracts on *E. coli* found that the tannins can auto-oxidize and form H_2_O_2_ (58). Tannin extracts which were prepared in oxic conditions inhibited the growth of *E. coli* in anoxic condition, but tannin extracts prepared anoxically were not inhibitory (58). Interestingly, this study also found that an *E. coli* strain constitutively expressing *oxyR* was more tolerant of the tannin extract compared to its parent strain (58). Although not yet investigated, tannins could be present in the AFEX hydrolysate and cause H_2_O_2_ to form. A chemical analysis found that switchgrass AFEX hydrolysate contains 2.3 µM *o*-methylgallic acid, a metabolite of gallic acid, as well as less than 1 µM gallic acid (2). Gallic acid is a main component of gallotannins. This suggests that tannin polymers may exist in hydrolysate and additional research is required to explore this hypothesis.

This study has important implications for the use of *Z. mobilis* as a biofuel producer. Although our fermentation experiments were conducted in an anaerobic chamber, in a real-world industrial setting, hydrolysate will be exposed to oxygen when it is produced and during the inoculation process. We observed that it took eight consecutive days of anoxic conditions for the peroxide concentrations in a 5 mL sample of AFEX hydrolysate to dissipate. In an industrial bioreactor, there will be a much larger volume of hydrolysate and it would be infeasible for it to become completely anoxic before starting the fermentation. This means that *Z. mobilis* will likely be exposed to even higher levels of peroxides in an industrial setting than what was observed in this study, underscoring the importance of mitigating oxidative stress for scale up. Further work is needed to understand the causes of peroxide formation in AFEX hydrolysate to determine whether it could be limited through optimized deconstruction methods. While overexpressing *oxyR* did not improve growth in hydrolysate, other genetic engineering strategies could be employed in *Z. mobilis* to better mitigate oxidative stress in large-scale fermentations. This could improve bacterial bioreactor performance and help advance the microbial biofuel industry.

## Materials and Methods

### Media and Chemicals

*Z. mobilis* strains were grown in *Zymomonas* rich medium (ZRMG), which contains 1% yeast extract, 2% D-glucose and 15 mM KH_2_PO_4_. Ammonia fiber expansion-treated hydrolysate (7% Glucan AFEX Switchgrass Hydrolysate (ASGH) and SynHv4.1 was from Great Lakes Bioenergy Research Center, Madison, Wisconsin (2). Hydrolysate pH was adjusted to 5.8 with 10 M NaOH. The hydrolysate was then sterilized by filtration. The components of SynHv4.1 can be found in Table S1. Anoxic experiments were conducted in an anaerobic chamber (Coy Laboratories) with an atmosphere containing ∼2% H_2_ balance N_2_. Media and supplies that were used for anoxic experiments were degassed for at least two days in the Coy Chamber. 100 µg/mL spectinomycin was added to cultures of *Z. mobilis* strains containing pRL814 plasmids. Catalase from bovine liver was from Sigma-Aldrich. Restriction enzymes, Q5 High-Fidelity Polymerase, and HiFi DNA Assembly Master Mix were from New England Biolabs. Oligonucleotides were from IDT.

### Bacterial Strains and Plasmids

Bacterial strains and plasmids are listed in Table 1. The broad host plasmid pRL814 is from Dr. Robert Landick (University of Wisconsin). The deletion strain was constructed as described previously (16). The plasmid pRL814*-oxyR* was constructed by first amplifying *oxyR* from the *Z. mobilis* genome, with primers that added overlapping sequences to pRL814 at the 5’ and 3’ ends. The fragment was then introduced to NdeI/BamHI digested pRL814 by Gibson Assembly. The construct was first transformed to *E. coli* Mach 1, and colony PCR was used to confirm insertion. Sanger sequencing was used to confirm the construct was correct. The donor strain *E. coli* WM6026 was used to conjugate this vector into ZM4 and *ΔoxyR*, as described in Lal et al (59).

**Table 1:** Strains and plasmids used in study.

| Strain or Plasmid | Description | Reference/Source |
| --- | --- | --- |
| **Strain** |  |  |
| *E. coli* Mach-1 | *E. coli* K-12 with genotype W $\Delta recA1398 endA1$ *fhuA* $\Phi 80\Delta(lac)M15 \Delta(lac)X74$ *hsdR* ($r_K^-m_K^+$) used for cloning constructs | ThermoFisher Scientific |
| *E. coli* WM6026 | Diaminopimelic acid (DAP) auxotroph donor strain used for conjugation of plasmids to *Z. mobilis* | (60) |
| ATCC 31821 (ZM4) | Wild-type *Z. mobilis* | Type Culture Collection |
| $\Delta oxyR$ | ZM4 with gene deletion of *oxyR* (ZMO1733) | This work |
| **Plasmids** |  |  |
| *pRL814* | *gfp* insert from *Aequorea victoria* | (61) |
| *pRL814-oxyR* | *oxyR* insert from *Z. mobilis* | This work |

### RNA Extraction and Sequencing

*Z. mobilis* strains were grown in oxic or anoxic conditions in ZRMG to an OD_600_ of approximately 1.0-2.0 and the culture volume that yielded 2 x 10^8^ cells was collected (for *Z. mobilis* ZM4: 6.55 x 10^7^ CFU/OD=1). The Qiagen RNeasy Mini Kit was used for RNA extraction. RNA quality was assessed by agarose gel and quantified using RNA fluorescence kit. RNA samples were sent to the Joint Genome Institute for Illumina RNA-sequencing.

### Transcriptomics Analysis

RNA-seq reads were processed with Trimmomatic version 0.39 and mapped to the *Zymomonas mobilis* ZM4 genome, GCF_003054575.1-RF_2023_03_19 genome using Bowtie 2 version 2.5.1 with default settings (62, 63). Read counts were calculated with HTSeq version 2.0.3 using the Z. *mobilis* ZM4 gene annotations (64). All raw data were deposited in the NCBI GEO database under project number GSE332662. edgeR version 4.0.16 was used for differential expression analysis.

### Bacterial Growth Curves

Strains were grown in ZRMG in an anaerobic chamber for 24 hours. Cultures were then diluted to starting OD_600_ of 0.1 in ZRMG or other media as indicated. Periodically, samples were taken for cell density measurements and HPLC analysis. Aerobic growth included shaking at 225 rpm at ambient oxygen condition. Anaerobic plate reader growth curves were conducted in a BioTek Synergy HTX multi-mode reader in the Coy chamber at 30°C, with orbital shaking at 282 cpm. Reads at optical density of 600 nm were taken every 15 minutes.

### HPLC Analysis

Samples were collected during growth curves and frozen at -20°C for storage. Samples were thawed before HPLC analysis and centrifuged at maximum speed for 10 minutes. Supernatants were transferred to glass vials for analysis. HPLC was performed on a Shimadzu 20A HPLC equipped with an Aminex HPX-87H, 300.0 × 7.8-mm column, at 60°C. Metabolites were eluted with 5 mM sulfuric acid at 0.6 mL/min, and a Refractive Index Detector was used for detection. Metabolite standards were run alongside samples for identification and quantification using external standard curves.

### Intracellular ROS Measurement

Bacterial cultures with a starting OD_600_ of 0.1 were grown for 24 hours (48 hours when grown in hydrolysate). Cultures were centrifuged and resuspended in Phosphate Buffered Saline (PBS) to an OD_600_ of 1. 200 µL of culture was added to a black 96-well plate. 25 µM CellRox green was then added to the samples. 100 µM Menadione was added 20 minutes prior to CellROX addition for positive control samples. The plate was shielded from light and incubated at 30°C for 30 minutes. Samples were collected from the plate and centrifuged at 8,000 rpm for 2 minutes. Samples were washed with 200 µL of PBS and transferred to new wells in the black 96-well plate. A BioTek Synergy H1 microplate reader was used to read fluorescence at 485 nm excitation, 520 nm emission.

### Peroxide Measurement

The Pierce Quantitative Peroxide Assay Kit was used to assess peroxide levels in the hydrolysates as described in the product manual. A BioTek Synergy H1 microplate reader was used to measure absorbance at 595 nm. Standards of 1 to 1,000 µM H_2_O_2_ were used to generate a standard curve. The linear portion of this standard curve was used to calculate the peroxide concentrations in the unknown samples.

## Acknowledgements

This material is based upon work at the Great Lakes Bioenergy Research Center supported by the U.S. Department of Energy, Office of Science, Biological and Environmental Research under Award Number DE-SC0018409. The work 509760 conducted by the U.S. Department of Energy Joint Genome Institute (https://ror.org/04xm1d337), a DOE Office of Science User Facility, is supported by the Office of Science of the U.S. Department of Energy under Contract No. DE-AC02-05CH11231.

## References

1. Zhou CH, Xia X, Lin CX, Tong DS, Beltramini J. 2011. Catalytic conversion of lignocellulosic biomass to fine chemicals and fuels. Chem Soc Rev 40:5588–5617.

2. Zhang Y, Serate J, Xie D, Gajbhiye S, Kulzer P, Sanford G, Russell JD, McGee M, Foster C, Coon JJ, Landick R, Sato TK. 2020. Production of hydrolysates from unmilled AFEX-pretreated switchgrass and comparative fermentation with Zymomonas mobilis. Bioresour Technol Rep 11:100517.

3. Rogers PL, Lee KJ, Skotnicki ML, Tribe DE. 1982. Ethanol Production by Zymomonas mobilis. Microbial reactions 1:37–84.

4. Bringer S, Finn RK, Sahm H. 1984. Effect of oxygen on the metabolism of Zymomonas mobilis. Arch Microbiol 139:376–381.

5. Felczak MM, TerAvest MA. 2022. Zymomonas mobilis ZM4 Utilizes an NADP+Dependent Acetaldehyde Dehydrogenase to Produce Acetate. J Bacteriol 204.

6. Yang S, Tschaplinski TJ, Engle NL, Carroll SL, Martin SL, Davison BH, Palumbo A V., Rodriguez M, Brown SD. 2009. Transcriptomic and metabolomic profiling of Zymomonas mobilis during aerobic and anaerobic fermentations. BMC Genomics 10.

7. Hayashi T, Furuta Y, Furukawa K. 2011. Respiration-deficient mutants of Zymomonas mobilis show improved growth and ethanol fermentation under aerobic and high temperature conditions. J Biosci Bioeng 111:414–419.

8. Kalnenieks U, Galinina N, Bringer-Meyer S, Poole RK. 1998. Membrane d-lactate oxidase in Zymomonas mobilis : evidence for a branched respiratory chain. FEMS Microbiol Lett 168:91–97.

9. Strohdeicher M, Neub B, Bringer-Meyer S, Sahm H. 1990. Electron transport chain of Zymomonas mobilis: Interaction with the membrane-bound glucose dehydrogenase and identification of ubiquinone 10. Arch Microbiol 154:536–543.

10. Kalnenieks U, Galinina N, Toma M, SkÄrds I. 2006. Electron transport chain in aerobically cultivated Zymomonas mobilis. FEMS Microbiol Lett 143:185–189.

11. Kalnenieks U, Galinina N, Irbe I, Toma M. 1995. Energy coupling sites in the electron transport chain of Zymomonas mobilis. FEMS Microbiol Lett 133:99–104.

12. Kainenieks U, De Graaf AA, Bringer-Meyer S, Sahm H. 1993. Oxidative phosphorylation in Zymomonas mobilis. Arch Microbiol 160:74–79.

13. Balodite E, Strazdina I, Galinina N, McLean S, Rutkis R, Poole RK, Kalnenieks U. 2014. Structure of the Zymomonas mobilis respiratory chain: Oxygen affinity of electron transport and the role of cytochrome c peroxidase. Microbiology (United Kingdom) 160:2045–2052.

14. Martien JI, Hebert AS, Stevenson DM, Regner MR, Khana DB, Coon JJ, Amador-Noguez D. 2019. Systems-Level Analysis of Oxygen Exposure in Zymomonas mobilis: Implications for Isoprenoid Production.

15. Enright AL, Banta AB, Ward RD, Vazquez JR, Felczak MM, Wolfe MB, TerAvest MA, Amador-Noguez D, Peters JM. 2023. The genetics of aerotolerant growth in an alphaproteobacterium with a naturally reduced genome. mBio 14.

16. Felczak MM, Bernard MP, TerAvest MA. 2023. Respiration is essential for aerobic growth of Zymomonas mobilis ZM4. mBio 14.

17. Demeester W, De Paepe B, De Mey M. 2024. Fundamentals and Exceptions of the LysR-type Transcriptional Regulators. ACS Synth Biol 13:3069–3092.

18. Chiang SM, Schellhorn HE. 2012. Regulators of oxidative stress response genes in Escherichia coli and their functional conservation in bacteria. Arch Biochem Biophys 10.1016/j.abb.2012.02.007.

19. Ming Zheng, Fredrik Aslund, Gisela Storz. 1998. Activation of the OxyR Transcription Factor by Reversible Disulfide Bond Formation. Science (1979) 279.

20. Toledano MB, Kullik I, Trinh F, Baird PT, Schneider TD, Storz G. 1994. Redox-dependent shift of OxyR-DNA contacts along an extended DNA-binding site: A mechanism for differential promoter selection. Cell 78:897–909.

21. Tao K, Fujita N, Ishihama A. 1993. Involvement of the RNA polymerase α subunit C-terminal region in co-operative interaction and transcriptional activation with OxyR protein. Mol Microbiol 7:859–864.

22. Wan F, Kong L, Gao H. 2018. Defining the binding determinants of Shewanella oneidensis OxyR: Implications for the link between the contracted OxyR regulon and adaptation. Journal of Biological Chemistry 293:4085–4096.

23. Silva LG, Lorenzetti APR, Ribeiro RA, Alves IR, Leaden L, Galhardo RS, Koide T, Marques M V. 2019. OxyR and the hydrogen peroxide stress response in Caulobacter crescentus. Gene 700:70–84.

24. Seib KL, Wu HJ, Srikhanta YN, Edwards JL, Falsetta ML, Hamilton AJ, Maguire TL, Grimmond SM, Apicella MA, McEwan AG, Jennings MP. 2007. Characterization of the OxyR regulon of Neisseria gonorrhoeae. Mol Microbiol 63:54–68.

25. Borisov VB, Siletsky SA, Nastasi MR, Forte E. 2021. ROS defense systems and terminal oxidases in bacteria. Antioxidants 10.

26. Choi H, Yang Z, Weisshaar JC. 2015. Single-cell, real-time detection of oxidative stress induced in Escherichia coli by the antimicrobial peptide CM15. Proc Natl Acad Sci U S A 112:E303–E310.

27. Chiou T-J, Tzeng W-F. 2000. The roles of glutathione and antioxidant enzymes in menadione-induced oxidative stress. Toxicology 154:75–84.

28. Myers KS, Park DM, Beauchene NA, Kiley PJ. 2015. Defining bacterial regulons using ChIP-seq. Methods. Academic Press Inc. 10.1016/j.ymeth.2015.05.022.

29. Tan T, Liu C, Liu L, Zhang K, Zou S, Hong J, Zhang M. 2013. Hydrogen sulfide formation as well as ethanol production in different media by cysND- and/or cysIJ-Inactivated mutant strains of Zymomonas mobilis ZM4. Bioprocess Biosyst Eng 36:1363–1373.

30. Serate J, Xie D, Pohlmann E, Donald C, Shabani M, Hinchman L, Higbee A, McGee M, La Reau A, Klinger GE, Li S, Myers CL, Boone C, Bates DM, Cavalier D, Eilert D, Oates LG, Sanford G, Sato TK, Dale B, Landick R, Piotrowski J, Ong RG, Zhang Y. 2015. Controlling microbial contamination during hydrolysis of AFEX-pretreated corn stover and switchgrass: Effects on hydrolysate composition, microbial response and fermentation. Biotechnol Biofuels 8.

31. Balan V, Bals B, Chundawat SPS, Marshall D, Dale BE. 2009. Lignocellulosic biomass pretreatment using AFEX. Methods Mol Biol 10.1007/978-1-60761-214-8_5.

32. Chundawat SPS, Venkatesh B, Dale BE. 2007. Effect of particle size based separation of milled corn stover on AFEX pretreatment and enzymatic digestibility. Biotechnol Bioeng 96:219–231.

33. Chundawat SPS, Vismeh R, Sharma LN, Humpula JF, da Costa Sousa L, Chambliss CK, Jones AD, Balan V, Dale BE. 2010. Multifaceted characterization of cell wall decomposition products formed during ammonia fiber expansion (AFEX) and dilute acid based pretreatments. Bioresour Technol 101:8429–8438.

34. Eckmann JB, Enright Steinberger AL, Davies M, Whelan E, Myers KS, Robinson ML, Banta AB, Lal PB, Coon JJ, Sato TK, Kiley PJ, Peters JM. 2025. Orthogonal chemical genomics approaches reveal genomic targets for increasing anaerobic chemical tolerance in Zymomonas mobilis. mSystems 10.1128/msystems.01001-25.

35. Barten LM, Crandall JG, Xie D, Serate J, Handowski E, Jen A, Overmyer KA, Coon JJ, Hittinger CT, Landick R, Zhang Y, Sato TK. 2025. pH adjustment increases biofuel production from inhibitory switchgrass hydrolysates. Bioresour Technol 432.

36. Simon P. Wolff. 1994. Ferrous Ion Oxidation in Presence of Ferric Ion Indicator Xylenol Orange for Measurement of Hydroperoxides. Methods Enzymol.

37. Li B, Liu N, Zhao X. 2022. Response mechanisms of Saccharomyces cerevisiae to the stress factors present in lignocellulose hydrolysate and strategies for constructing robust strains. Biotechnology for Biofuels and Bioproducts. BioMed Central Ltd 10.1186/s13068-022-02127-9.

38. Geng B, Liu S, Chen Y, Wu Y, Wang Y, Zhou X, Li H, Li M, Yang S. 2022. A plasmid-free Zymomonas mobilis mutant strain reducing reactive oxygen species for efficient bioethanol production using industrial effluent of xylose mother liquor. Front Bioeng Biotechnol 10.

39. Wang Y, Geng B, Liu S, Wu Y, Yang S. 2026. S267P mutation of OxyR regulator in Zymomonas mobilis: mechanism of oxidative stress tolerance and applications in cellulosic hydrolysate fermentation and oxidative stress monitoring. Synth Syst Biotechnol 13:531–544.

40. Yang Q, Yang Y, Tang Y, Wang X, Chen Y, Shen W, Zhan Y, Gao J, Wu B, He M, Chen S, Yang S. 2020. Development and characterization of acidic-pH-tolerant mutants of Zymomonas mobilis through adaptation and next-generation sequencing-based genome resequencing and RNA-Seq. Biotechnol Biofuels 13.

41. Seaver LC, Imlay JA. 2001. Alkyl hydroperoxide reductase is the primary scavenger of endogenous hydrogen peroxide in Escherichia coli. J Bacteriol 183:7173–7181.

42. Poole LB. 2005. Bacterial defenses against oxidants: Mechanistic features of cysteine-based peroxidases and their flavoprotein reductases. Arch Biochem Biophys 10.1016/j.abb.2004.09.006.

43. Gonzalez-Flechat B, Demple B. 1995. Metabolic Sources of Hydrogen Peroxide in Aerobically Growing Escherichia coli. J Biol Chem 270:13681–13687.

44. Mironov A, Seregina T, Nagornykh M, Luhachack LG, Korolkova N, Lopes LE, Kotova V, Zavilgelsky G, Shakulov R, Shatalin K, Nudler E. 2017. Mechanism of H2S-mediated protection against oxidative stress in Escherichia coli. Proc Natl Acad Sci U S A 114:6022–6027.

45. Konstantin Shatalin, Elena Shatalina, Alexander Mironov, Evgeny Nudler. 2011. H2S: A Universal Defense Against Antibiotics in Bacteria. Science (1979) 334:986–989.

46. Tikhomirova A, Rahman MM, Kidd SP, Fererro RL, Roujeinikova A. 2024. Cysteine and resistance to oxidative stress: implications for virulence and antibiotic resistance. Trends Microbiol. Elsevier Ltd 10.1016/j.tim.2023.06.010.

47. Lluis Masip, Karthik Veeravalli, George Georgiou. 2006. The many faces of glutathione in bacteria. Antioxid Redox Signal 8.

48. Wang J, Liu Jun, Zhao Yuqiang, Sun Minghui, Yu Guixi, Fan Jiaqin, Tian Yanli, Hu Baishi. 2022. OxyR contributes to virulence of Acidovorax citrulli by regulating anti-oxidative stress and expression of flagellin FliC and type IV pili PilA. Front Microbiol.

49. Hennequin C, Forestier C. 2009. oxyR, a LysR-type regulator involved in Klebsiella pneumoniae mucosal and abiotic colonization. Infect Immun 77:5449–5457.

50. Shanks RMQ, Stella NA, Kalivoda EJ, Doe MR, O’Dee DM, Lathrop KL, Feng LG, Nau GJ. 2007. A Serratia marcescens OxyR homolog mediates surface attachment and biofilm formation. J Bacteriol 189:7262–7272.

51. Chung CH, Fen SY, Yu SC, Wong HC. 2016. Influence of oxyR on growth, biofilm formation, and mobility of Vibrio parahaemolyticus. Appl Environ Microbiol 82:788–796.

52. Sun W, Hattman S. 1996. Escherichia coli OxyR protein represses the unmethylated bacteriophage Mu mom operon without blocking binding of the transcriptional activator C. Nucleic Acids Res 24:4042–4049.

53. Glinkowska M, Łoś JM, Szambowska A, Czyż A, Całkiewicz J, Herman-Antosiewicz A, Wróbel B, Wȩgrzyn G, Węgrzyn A, Łoś M. 2010. Influence of the Escherichia coli oxyR gene function on λ prophage maintenance. Arch Microbiol 192:673–683.

54. Yordi EG, Molina Pérez E, Matos MJ, Villares EU. Antioxidant and Pro-Oxidant Effects of Polyphenolic Compounds and Structure-Activity Relationship Evidence.

55. Babich H, Schuck AG, Weisburg JH, Zuckerbraun HL. 2011. Research strategies in the study of the pro-oxidant nature of polyphenol nutraceuticals. J Toxicol 10.1155/2011/467305.

56. Kuzina SI, Shilova IA, Mikhailov AI. 2011. Chemical and radiation-chemical radical reactions in lignocellulose materials. Radiation Physics and Chemistry 80:937–946.

57. Hassanp S, Maheri-Sis N, Eshratkhah B, Baghbani F, Hassanpour S, Mehmandar FB. 2011. Plants and secondary metabolites (Tannins): A Review. International Journal of Forest, Soil and Erosion 1:47–53.

58. Smith AH, Imlay JA, Mackie RI. 2003. Increasing the oxidative stress response allows Escherichia coli to overcome inhibitory effects of condensed tannins. Appl Environ Microbiol 69:3406–3411.

59. Lal PB, Wells FM, Lyu Y, Ghosh IN, Landick R, Kiley PJ. 2019. A Markerless Method for Genome Engineering in Zymomonas mobilis ZM4. Front Microbiol 10:1–11.

60. Blodgett JAV, Thomas PM, Li G, Velasquez JE, Van Der Donk WA, Kelleher NL, Metcalf WW. 2007. Unusual transformations in the biosynthesis of the antibiotic phosphinothricin tripeptide. Nat Chem Biol 3:480–485.

61. Ghosh IN, Martien J, Hebert AS, Zhang Y, Coon JJ, Amador-Noguez D, Landick R. 2019. OptSSeq explores enzyme expression and function landscapes to maximize isobutanol production rate. Metab Eng 52:324–340.

62. Bolger AM, Lohse M, Usadel B. 2014. Trimmomatic: A flexible trimmer for Illumina sequence data. Bioinformatics 30:2114–2120.

63. Langmead B, Salzberg SL. 2012. Fast gapped-read alignment with Bowtie 2. Nat Methods 9:357–359.

64. Putri GH, Anders S, Pyl PT, Pimanda JE, Zanini F. 2022. Analysing high-throughput sequencing data in Python with HTSeq 2.0. Bioinformatics 38:2943–2945.

